# Evaluation of the GenoType MTBDR*plus* and MTBDR*sl* for the detection of drug-resistant *Mycobacterium tuberculosis* on isolates from Beijing, China

**DOI:** 10.1101/311944

**Authors:** Jiyong Jian, Xinyu Yang, Jun Yang, Liang Chen

**Author notes:** Correspondence to: Dr Liang Chen Address: Department of Clinical Laboratory, Beijing Shijitan Hospital, Capital Medical University, NO. 10, Tie Yi Road, Yang Fang Dian, Haidian District, Beijing, 100038, China.

## Abstract

The incidence of tuberculosis (TB) and especially multidrug-resistant TB (MDR) and extreme drug resistance (XDR-TB) continue to increase alarmingly worldwide and reliable and fast diagnosis of MDR-TB and XDR-TB is essential for the adequate treatment of patients. So molecular line probe assays (LPAs) for detection of MDR-TB and XDR-TB have been endorsed by the World Health Organization (WHO). We analyzed 96 isolates from Beijing comparing culture-based drug susceptibility testing (DST) to LPAs detecting rifampicin (RFP), isoniazid (INH), and levofloxacin (LFX), amikacin (AM), capreomycin (CMP), ethambutol (EMB) resistance. Compared to phenotypic DST, the GenoType MTBDR*plus* and MTBDR*sl* showed a sensitivity of 98.7% and a specificity of 88.9% for detection of RFP resistance, 82.1% and 94.4% for INH, 89.7% and 94.4% for LFX, 60.0% and 98.7% for AM/CPM, 57.5% and 98.2% for EMB, respectively. The sensitivity and specificity of LPAs for MDR-TB and XDR-TB were 80.8%, 100% and 50.0%, 97.6%. Mutations in codon S531L of the *rpoB* gene and S315T1 of *KatG* gene were dominated in MDR-TB strains. The most frequently observed mutations were in codon A90V of the *gyrA* gene, A1401G of the *rrs* gene and M306V of the *embB* gene, according to the MTBDR*sl* results. Our study showed that, in combination to phenotypic DST, application of the LPAs might be an efficient and reliable supplementary DST assay for rapid susceptibility screening of MDR-TB and XDR-TB. Using LPAs in high MDR/XDR burden countries allows for appropriate and timely treatment, which will reduce transmission rates, morbidity and improve treatment outcomes in patients.

## INTRODUCTION

Tuberculosis (TB) remains a global public health threat, especially in developing countries. According to the global TB report of the World Health Organization (WHO), in 2016, TB was responsible for the deaths of 1.67 million people and generated 10.4 million cases, of which 490,000 considered as multidrug resistant (MDR-TB), and 8014 showed extreme drug resistance (XDR-TB) (1). In China, a high TB-burden country, approximately110,000 new MDR-TB and 8,200 new XDR-TB cases emerge yearly according to a national survey held in 2007(2). According to another report, about 55% of MDR-TB cases in China have not been identified (3). MDR-TB is TB that is resistant to both rifampicin (RFP)and isoniazid (INH), the two most powerful anti-TB drugs; it requires treatment with a second-line regimen. XDR-TB is defined as MDR-TB plus resistance to at least one fluoroquinolone (FQ) and a second line injectable drug (SLID), including amikacin (AM), capreomycin (CPM) and kanamycin (KM), the two most important classes of medicines in an MDR-TB regimen.

Detection and treatment of MDR-TB and XDR-TB requires susceptibility testing to screen for resistance to specific antibiotics and utilizing results to design a treatment regimen. Conventional culture-based phenotypic drug susceptibility testing (DST) methods, considered to be the gold standard for drug resistance determination, is important for MDR-TB confirmation and the assessment of drug resistance to second line and new drugs in the management of MDR-TB and XDR-TB. But phenotypic DST methods can take months to get results(4). Molecular-based assays designed to detect specific drug resistance-encoding mutations in *M. tuberculosis(MTB)* have the advantage of achieving faster resistance results within 48 hours compared to conventional DST. In 2008, WHO endorsed the use of the molecular test GenoType® MTBDR*plus* (Hain Lifescience, Nehren, Germany) for rapid detection of resistance to RFP and INH. The assay detects mutations in the *rpoB* gene for RFP resistance, in the *katG* gene for high-level INH resistance and in the *inhA* regulatory region gene for low-level INH resistance (5).In May 2016, the WHO recommended use of GenoType® MTBDR*sl* (Hain Lifescience, Nehren, Germany) to detect mutations in the *gyrA, rrs*, and *embB* genes and, therefore, resistance to FQ, SLID and ethambutol (EMB) allowing diagnosis of XDR-TB among MDR-TB patients (6).

In order to rapidly detect MDR-TB and XDR-TB, WHO endorsed line probe assays (LPA) of MTBDRplus and MTBDRsl for the detection of MTB, RFP, INH, FQ and SLID resistance in acid-fast bacilli smear positive sputum or MTB cultures in 2017.The aim of this study was to compare the diagnostic performance of the MTBDRplus and MTBDRsl assays for the detection of MDR-TB and XDR-TB to the gold standard phenotypic DST, using culture isolates obtained from patients in Beijing.

## MATERIALS AND METHODS

### Study design

The evaluation of the GenoType MTBDR*plus*v1.0and MTBDR*sl* v1.0assays was conducted at the Beijing Shijitan Hospital and Beijing research institute for tuberculosis control. The study was approved by the Ethics Committee of Beijing Shijitan Hospital, Beijing research institute for tuberculosis control and PLA 306 Hospital. A total of 96 MTB isolates were collected by the Beijing research institute for tuberculosis control from Beijing between 2015 to 2016, including 78 MD-TB, 12 XDR-TB and 18 randomly chosen fully susceptible isolates based on phenotypic DST. We compared the performance of the MTBDR*plus*v1.0 and MTBDR*sl* v1.0 assays to phenotypic MTB DST for susceptibility testing of first and second-line anti-TB drugs.

### Drug susceptibility testing

Phenotypic DST was performed against RFP, INH, Levofloxacin(LFX), AM, CPM and EMB using the standard version of the WHO proportion method on Lowenstein-Jensen medium (L-J)(7,8) and considered the gold standard for resistance determination. The following critical concentrations of drugs recommended by WHO for testing of drug-resistant TB using proportion method DST were used: RFP 40μg/ml,INH 0.2μg/ml, LFX 2μg/ml, AM 30μg/ml, CPM 40μg/ml and EMB 2μg/ml. Growth on the control medium was compared with growth on a drug-containing medium to determine susceptibility.

### Line probe assays

The MTBDR*plus* and the MTBDR*sl* assays were performed directly on the MTB isolates according to the manufacturer’s instructions. The person performing the tests was blinded to the proportion method DST. In order to give a valid result, all six expected control bands should appear correctly. Otherwise, the result is considered invalid. The absence of at least one of the wild-type bands or the presence of bands indicating a mutation in each drug resistance-related gene implies that the sample tested is resistant to the respective antibiotic. When all the wild-type probes of a gene stain positive and there is no detectable mutations within the region examined, the sample tested is susceptible to the respective antibiotic.

### Quality control

The fully susceptible *M. tuberculosis* H37Rv reference strain was used as quality control (QC) for proportion method DST and LPA. This QC strain is susceptible to first- and second-line drugs tested in this study.

### Statistical analysis

Sensitivity, specificity, negative predictive value (NPV), positive predictive value (PPV) and agreement of LPA compared to proportion method DST were calculated. The precision of the estimates was reported using95% confidence intervals (95%CI).*P* ≤0.05 was considered statistically significant. Agreement between the two methods was assessed using the kappa statistic. All data were analyzed by SPSS version19.0 (SPSS Inc, Chicago, IL,USA).

## RESULTS

### Phenotypic DST results

Table 1 summarizes drug susceptibility patterns of isolates included in the study. An isolate collection (n=96) containing 78 MDR-TB (81.3%) and 12 MDR-TB isolates were XDR-TB (12.5%).Among these isolates, 78(81.3%)were resistant to RFP, 78(81.3%)were resistant to INH, 39 (40.6%) were resistant to LFX, 11 (11.5%) were resistant to AM, 18 (18.8%) were resistant to CM and 40 (41.7%) were resistant to EMB by conventional phenotypic DST.

**Table 1.** Phenotypic DST profiles of 96 isolates

| Drug resistance | No. of isolates(%) |
| --- | --- |
| MDR-TB | 78 (81.3) |
| XDR-TB | 12 (12.5) |
| Any resistance to |  |
| RFP | 78 (81.3) |
| INH | 78 (81.3) |
| LFX | 39 (40.6) |
| AM | 11 (11.5) |
| CPM | 18 (18.8) |
| EMB | 40 (41.7) |

### Performance of GenoType MTBDR*plus* and MTBDR*sl* assay

The performance of GenoType MTBDR*plus* was summarized in Table 2. The sensitivity for RFP resistance was determined as 98.7%(95%CI 96.2–100),specificity 88.9%(95%CI 72.8–100),PPV 97.5%(95%CI 93.9–100),NPV 94.1%(95%CI 81.6–100)and diagnostic efficacy96.9%; for INH resistance, sensitivity 82.1%(95%CI 73.3–90.8), specificity 94.4%(95%CI 82.7–100), PPV 98.5%(95%CI 95.4–100),NPV 54.8%(95%CI 36.3–73.4) and diagnostic efficacy84.4% and for MDR-TB, sensitivity 80.8%(95%CI 71.8–89.7),specificity 100%(95%CI 100–100), PPV 100%(95%CI 100–100),NPV 54.5%(95%CI 36.6–72.5)and diagnostic efficacy84.4%, compared to phenotypic DST.

**Table 2.** Performance of GenoType MTBDR*plus* and GenoType MTBDR*sl* assay compared to phenotypic DST

| Genotypic DST<br>results (n=96) | phenotypic | DST results | %Sensitivity | %Specificity | %PPV(95%CI) | %NPV(95%CI) | Diagnostic<br>efficacy(%) | K-Value |
| --- | --- | --- | --- | --- | --- | --- | --- | --- |
|  | Resistant | Sensitive | (95%CI) | (95%CI) |  |  |  |  |
| RFP |  |  |  |  |  |  |  |  |
| Resistant | 77 | 2 | 98.7(96.2–100) | 88.9(72.8–100) | 97.5(93.9–100) | 94.1(81.6–100) | 96.9 | 0.895 |
| Sensitive | 1 | 16 |  |  |  |  |  |  |
| INH |  |  |  |  |  |  |  |  |
| Resistant | 64 | 1 | 82.1(73.3–90.8) | 94.4(82.7–100) | 98.5(95.4–100) | 54.8(36.3–73.4) | 84.4 | 0.599 |
| Sensitive | 14 | 17 |  |  |  |  |  |  |
| MDR-TB |  |  |  |  |  |  |  |  |
| Resistant | 63 | 0 | 80.8(71.8–89.7) | 100(100–100) | 100(100–100) | 54.5(36.6–72.5) | 84.4 | 0.612 |
| Sensitive | 15 | 18 |  |  |  |  |  |  |
| LFX |  |  |  |  |  |  |  |  |
| Resistant | 35 | 2 | 89.7(79.8–99.7) | 96.5(91.6–100) | 94.6(87.0–100) | 93.2(86.6–99.8) | 93.8 | 0.869 |
| Sensitive | 4 | 55 |  |  |  |  |  |  |
| AM/CPM |  |  |  |  |  |  |  |  |
| Resistant | 12 | 1 |  |  |  |  |  |  |
|  |  |  | 60.0(36.5–83.5) | 98.7(96.1–100) | 92.3(75.5–100) | 90.4(83.9–96.8) | 90.6 | 0.674 |
| Sensitive | 8 | 75 |  |  |  |  |  |  |
| EMB |  |  |  |  |  |  |  |  |
| Resistant | 23 | 1 |  |  |  |  |  |  |
|  |  |  | 57.5(41.5–73.5) | 98.2(94.6–100) | 95.8(87.2–100) | 76.4(66.3–86.4) | 81.2 | 0.591 |
| Sensitive | 17 | 55 |  |  |  |  |  |  |
| XDR-TB |  |  |  |  |  |  |  |  |
| Resistant | 6 | 2 |  |  |  |  |  |  |
|  |  |  | 50.0(16.8–83.2) | 97.6(94.3–100) | 75.0(36.3–100) | 93.2(87.8–98.6) | 91.7 | 0.556 |
| Sensitive | 6 | 82 |  |  |  |  |  |  |

AK/CPM indicates phenotypic resistance to Amikacine and/or Capreomycin

The performance of GenoType MTBDR*sl* was also showed in Table 2. The sensitivity for detecting LFX resistance was 89.7% (95%CI79.8–99.7), specificity 96.5% (95%CI 91.6–100), PPV 94.6% (95%CI 87.0–100), NPV 93.2% (95%CI 86.6–99.8) and diagnostic efficacy 93.8%; for AM/CPM, sensitivity 60.0% (95%CI 36.5–83.5), specificity 98.7% (95%CI 96.1–100), PPV 92.3% (95%CI 75.5–100), NPV 90.4% (95%CI 83.9–96.8) and diagnostic efficacy90.6%; for EMB, sensitivity 57.5% (95%CI 41.5–73.5), specificity 98.2% (95%CI 94.6–100), PPV 95.8% (95%CI 87.2–100), NPV 76.4% (95%CI 66.3–86.4) and diagnostic efficacy81.2%; while for XDR-TB, sensitivity 50.0% (95%CI 16.8–83.2), specificity 97.6% (95%CI 94.3–100), PPV 75.0% (95%CI 36.3–100), NPV 93.2% (95%CI 87.8–98.6) and diagnostic efficacy 91.7%.

### Detection of mutations associated with drug resistance using MTBDR*plus* and MTBDR*sl* assay

The distribution of mutations is summarized in Table 3. Mutations in *rpoB* conferring resistance to RFP were detected in 82.3% (79/96) of the isolates. The RFP resistant isolates displayed different mutations: 58.2% (46/79) of the isolates had mutation at positionS531L, 10.1% (8/79) of the isolates had mutation at position D516V, 8.9% (7/79) of the isolates had mutation at position H526Y, while in fifteen isolates, and mutation was detected only at the wild type probes. Of the fifteen isolates with mutation that detected only at wild type probes, seven isolates had mutation atrpoBWT7, five isolates atWT8, two isolates atWT3 and WT4 and one isolate at WT2 and WT3. According to the kit manufacturer’s recommendation, these isolates were considered resistant and indicated the presence of a less common or rare mutation.

**Table 3.**
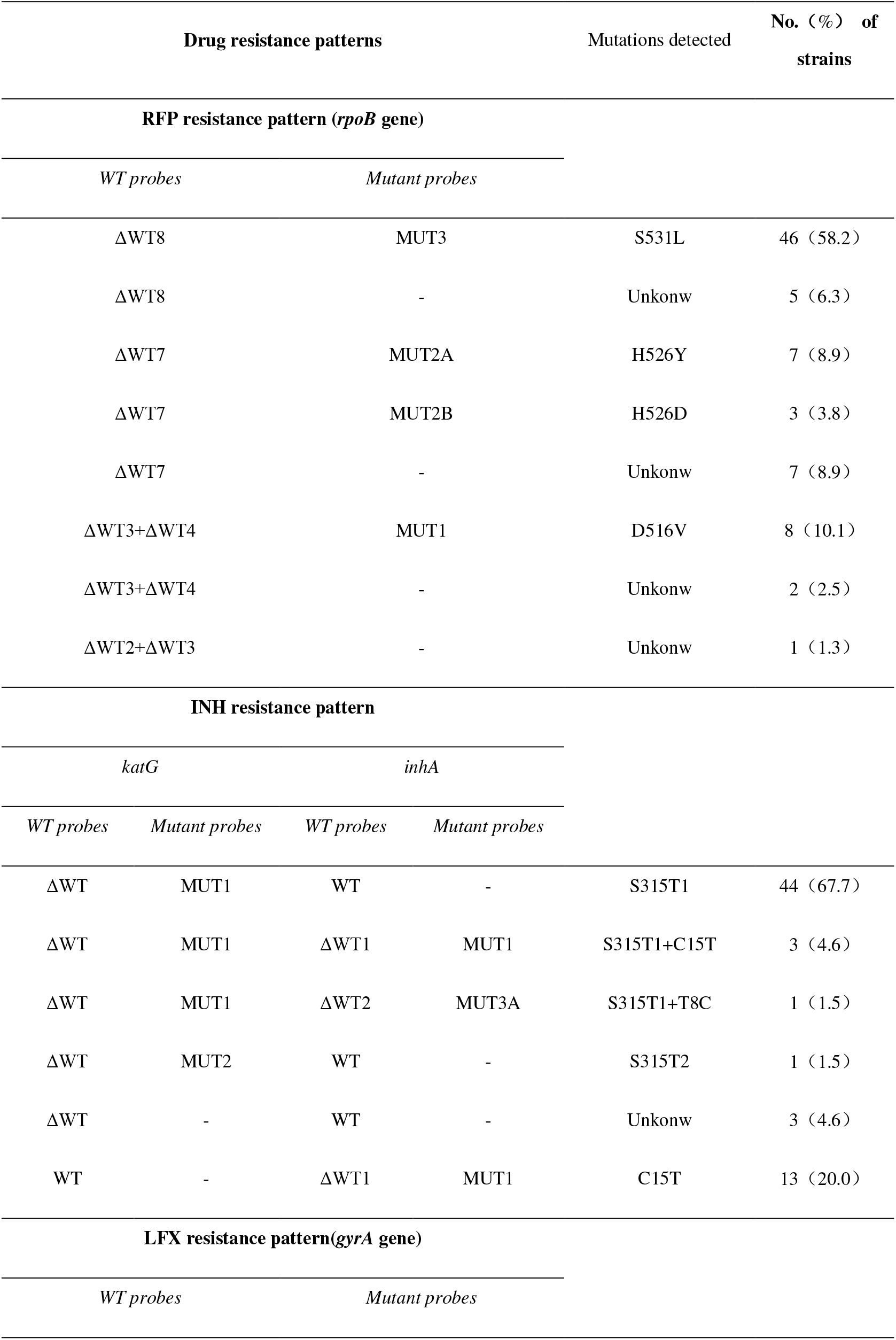
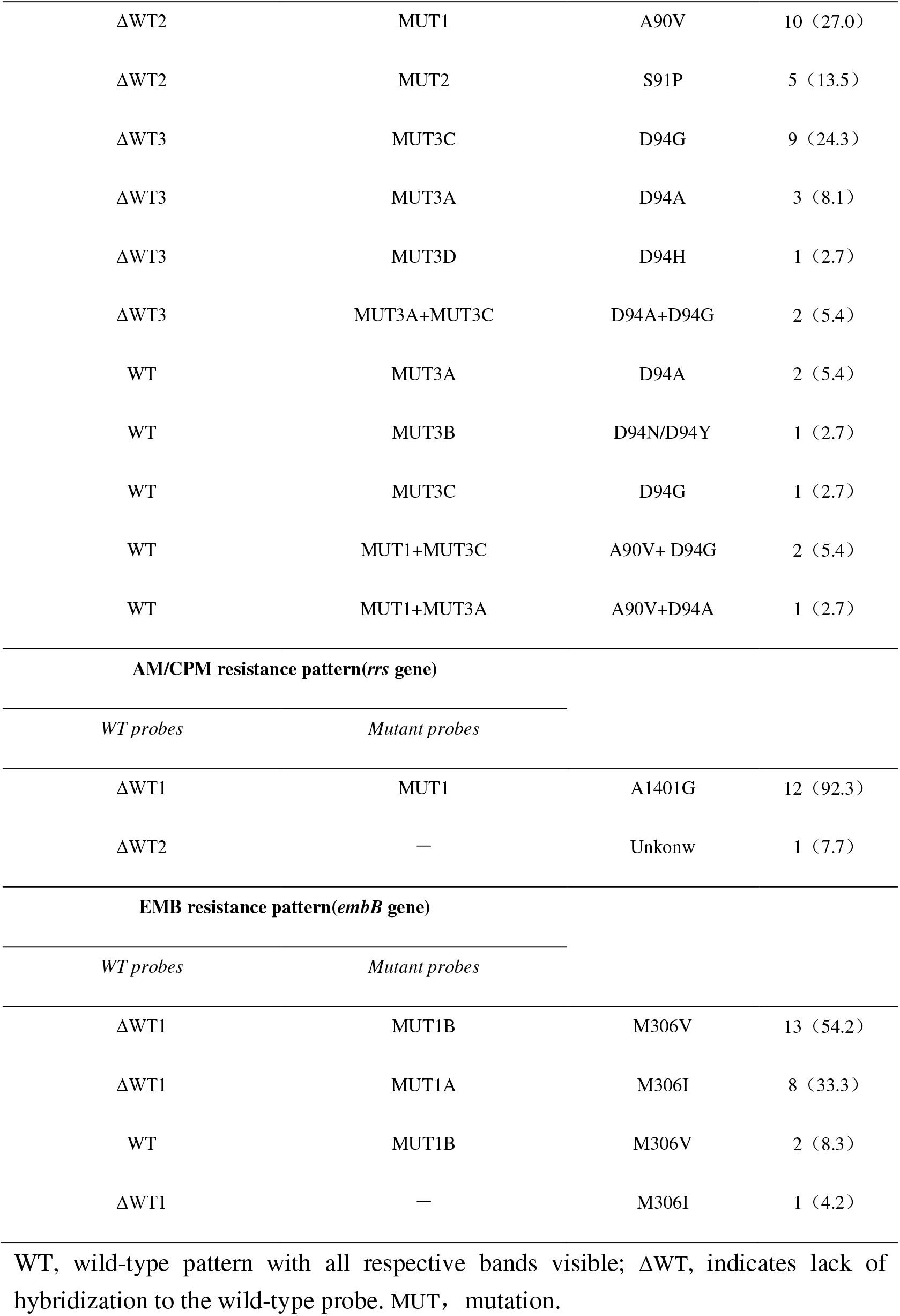
Variety of Drug resistance patterns using GenoType MTBDR*plus* and MTBDR*sl* assay

67.7% (65/96) of the isolates showed mutation in *katG* gene or *inhA* promoter region indicating that they were resistant to INH. There were four isolates that showed mutations at both *katG* and *inhA* gene. Of 65 INH resistant strains, 73.8% (48/65) had mutation in the *katG* gene with amino acid change ofS315T1, indicating high level resistance, while 24.6 %(16/65) of the strains had mutation in the *inhA* gene, C15T, indicating low level resistance. One isolate with mutation can only detected at wild type probes.

There were 38.5% (37/96) isolates resistant to LFX as tested by LPA. The majority of the *gyrA* mutations, 37.8% (14/37), were observed at D94G. Other *gyrA* mutations detected by the assay were at A90V (13/37; 35.1%), at D94A (8/37; 21.6%) and at S91P (5/37; 13.5%). 59.5% (22/37) mutations in *gyrA* were detected at codon 94.

LPA detected 12 defined mutations and 1 undefined mutations in the *rrs* gene among the 13/96 (13.5%) isolates. The most frequently observed mutation (12/13; 92.3%) for AM/CPM resistance was *rrs* MUT1 (A1401G).

EMB resistance was detected in 24 (25.0%) of 96isolates, in which the mutation M306V was the most prevalent (15/24; 62.5%), followed by the mutation M306I (9/24; 37.5%) in *embB* gene.

## DISCUSSION

Drug resistant TB poses a great threat to TB control programs worldwide. Early diagnosis and effective treatment requires a sensitive and specific diagnostic tool. According to the WHO, LPA is an optimal susceptibility testing of MTB to anti-TB drugs for an effective treatment regimen (1).In this study, we evaluated the diagnostic accuracy of MTBDR*plus* and MTBDR*sl* using culture isolates in Beijing.

MTBDR*plus* showed high sensitivity and specificity for the detection of susceptibility to RFP (sensitivity-98.7% andspecificity-88.9%). The sensitivity for susceptibility to RFP was in concordance with the high sensitivity of the MTBDR*plus* assay in the range of 95–100%(9,10,11). In this study, the S531L mutation in *rpoB* was the most frequent (58.2%), followed by the D516Vmutation(10.1%). This is similar to the frequencies reported in other studies(11,12).This high rate of detection can be explained by the fact that the mutations responsible for RFP resistance are mainly located in the 81-bp hot-spot region and that mutations outside this location are rare and are associated with low-level resistance(13,14).

The sensitivity and specificity of MTBDR*plus* for the detection of susceptibility to INH were 82.1% and 94.4% respectively. In our study, the sensitivity for INH resistance was much lower than 86–100% (15,16) and higher than 69.9%, another study using clinical isolates in Eastern China (17).As for INH resistance, of the 65 resistant isolates that the Genotype MTBDR*plus* detected, 73.8% of them carried a mutation at theS315T1 codon of the *katG* gene and 24.6% with the mutation C15Tin the *inhA* regulatory region. As has previously been described by several authors, the most common mutation involved in INH resistance is the S315T substitution in *katG*, which has also been related to high levels of INH resistance (18,19).The most prevalent mutation in the *inhA* gene detected using LPA in our study was C15T with loss of WT1, which confers low-level INH resistance; this is also supported by previous studies (20,21). The relatively low sensitivity to detect INH resistance for MTBDR*plus* is due to the complex molecular basis, because it involves mutations in more than one gene or gene complex, such as the *katG, inhA*, and *kasA* genes and the intergenic region of the *oxyR-ahpC* complex (22,23).

Previous studies have shown that the sensitivity of GenoType MTBDR*sl* assay to be ranged from 75.6% to 90.6% for detecting FQ resistance (24,25,26). In this study, the sensitivity of the MTBDR*sl* assay for detecting LFX resistance in clinical strains was 89.7%. FQ resistance in MTB is ascribed mainly to *gyrA* mutations, with 57.5% of mutations detected at codon 94 and 31.5% at codon 90 (27). Consistent with published data, the highest frequency of mutations conferring FQ resistance was observed in *gyrA* codon 94(59.5%), followed by codon 90(13/37, 35.1%). Our study showed that the most prevalent mutation pattern in *gyrA* was the D94G mutation, followed by theA90V and D94A mutation, which is in agreement to other studies (24,28). Given that mutation probes for *gyrB* are not included in the assay, it is possible that some isolates might have had mutations in these positions, which the assay could not detect.

Our study showed a low sensitivity of the LPA for the detection of resistance to SLID (60.0%), which was much lower than other reports (86.7%,100%)(29,30) and was similar to Zeng’s report(28). The variable results may be ascribed to the fact that geographically different MTB lineages could result in different gene mutation patterns. Interestingly, cross-resistance to AM, KM, and CPM has been reported (31). The predominant *rrs* gene mutation was A1401G detected by MTBDR*sl* assay (92.3%) according to our study. Consistent with published data (30,32), *rrs* MUT1 A1401G was the most frequently observed mutation among tested isolates. In this study, we found 8 isolates which MTBDR*sl* assay showed sensitive to SLID, while the phenotypic DST indicated that they were resistant to CPM but sensitive to AM. Therrs1401 mutation alone was not found with sufficient frequency to detect more than 70–80% of global MTB strains resistant to AM and CPM, while the *eis* promoter, *tlyA* and *gidB* appeared to be involved in the resistance to AM and CPM (33).

The sensitivity of the MTBDR*sl* for detecting EMB resistance was 57–69.2%(24,25,26). Our results confirmed the poor performance of *embB* mutations in detecting EMB resistance. A recent meta-analysis showed a similar sensitivity with the MTBDR*sl* assay for detecting EMB resistance (55%) (34).The most common mutations detected in *embB* by the MTBDR*sl* assay were M306V in 62.5% and M306I in 37.5% of EMB-resistant isolates, corresponded to previous report (35). This suggests that the significance of mutations in this codon is a limited and there isa need to identify other mutations conferring resistance to thisdrug. Huang *et al*. identified several mutations in *embB* other than that at codon 306 (35).At the same time, recent results on the proficiency testing of DST in supranational TB reference laboratories highlighted a lack of consistency in DST results for EMB and the reproducibility was also found to be poor(36).

As a rapid diagnosis of MDR-TB or even XDR-TB is of a high importance for the patient outcome and of a high epidemiological importance, we evaluated the GenoType MTBDR*plus* and MTBDR*sl* assays on the MTB isolates. These tests allow information about the MTB resistance pattern within 1 day, and the conventional DST testing takes 2 weeks. The sensitivity (80.8%) for the detection of MDR-TB in the present study was much lower than previous report (10).We also found that GenoType MTBDR*sl* was specific (97.6%) for the diagnosis of XDR-TB, although the sensitivity is very low (50.0%), as reported in previous studies(17).

There are several limitations to our work. MTBDR*plus* version 1 and MTBDR*sl* version 1 were used, which have recently been succeeded by a new iteration (version 2) (37,38). The new MTBDR*plus* v2.0 test with a higher analytical sensitivity when compared with the original MTBDR*plus*, which allows this new version to be performed on both smear-positive and smear-negative clinical specimens. GenoType MTBDR*sl* VER 2.0 is redesigned based on GenoType MTBDR*sl* VER 1.0 and accommodates additional mutations for the molecular detection of resistance to FLQ involving *gyrA* and *gyrB* and SLID resistance covering both *rrs* and *eis* genes (39). A further limitation was our samples size, particularly of the second-line anti-tuberculosis drugs resistant group was small. Finally, the study was that both LPA assays were only tested on the MTB isolates and not patient’s samples. The time to be positive of MTB cultures can take up to 7 weeks, while rapid and safe diagnosis of MDR and XDR tuberculosis is essential for the adequate treatment of patients.

In conclusion, discordance still exists between conventional and LPA approaches in resistance testing. So the negative results require further investigation as resistance to second-line drugs may still be present but undetected by LPA assays. Even though MTBDR*sl* had suboptimal diagnostic sensitivity for FQ, SLID and EMB, MTBDR*plus* and MTBDR*sl* remain an important supplementary role for rapidly detect MDR-TB and XDR-TB, given that phenotypic DST has a prolonged within-laboratory turn-around-time and are technically challenging. We suggest that LPAs could be used as a supplementary method for the detection of MDR-TB and XDR-TB in high burden countries to improve treatment outcomes in patients. Reducing the time to diagnosis, commencing appropriate therapy timeously and preventing transmission of drug-resistant strains are major advantages of LPAs.

## ACKNOWLEDGMENTS

We thank all of the persons involved in this study for their essential work and support. We also wish to thank bioMérieux Shanghai Co. Limited, for their technical support for this study.

We have no conflicts of interest to declare.

## REFERENCE

1. World Health Organization. Global Tuberculosis Report. 2017th ed. World Health Organization; 2017.

2. Zhao Y1, Xu S, Wang L, Chin DP, Wang S, Jiang G, Xia H, Zhou Y, Li Q, Ou X, Pang Y, Song Y, Zhao B, Zhang H, He G, Guo J, Wang Y. 2012. National survey of drug-resistant tuberculosis in China. N Engl J Med 366:2161–2170.

3. World Health Organization. Global Tuberculosis Report. 2014th ed. World Health Organization; 2014.

4. Bwanga F, Hoffner S, Haile M, Joloba ML. 2009. Direct susceptibility testing for multi-drug resistant tuberculosis: A meta-analysis. BMC Infect Dis 9: 67.

5. Piersimoni C, Olivieri A, Benacchio L, Scarparo C. 2006.Current perspectives on drug susceptibility testing of Mycobacterium tuberculosis complex: the automated nonradiometric systems. J Clin Microbiol 44: 20–28.

6. Theron G, Peter J, Richardson M, Barnard M, Donegan S, Warren R, Steingart KR, Dheda K. 2014. The diagnostic accuracy of the Genotype® Mtbdrsl asssay for the detection of resistance to second-line anti-tuberculosis drugs. Cochrane Database Syst Rev 10:CD010705.

7. Canetti G, Fox W, Khomenko A, Mahler HT, Menon NK, Mitchison DA, Rist N, Smelev NA. 1969. Advances in techniques of testing mycobacterial drug sensitivity, and the use of sensitivity tests in tuberculosis control programmes. Bull World HealthOrgan 41:21–43.

8. Canetti G, Froman S, Grosset J, Hauduroy P, Langerova M, Mahler HT, Meissner G, Mitchison DA, Sula L. 1963. Mycobacteria: laboratory methods for testing drug sensitivity and resistance. Bull World Health Organ 29:565–78.

9. Asante-Poku A, Otchere ID, Danso E, Mensah DD, Bonsu F, Gagneux S, Yeboah-Manu D. 2015. Evaluation of GenoType MTBDRplus for the rapid detection of drug-resistant tuberculosis in Ghana. Int J Tuberc LungDis 19:954–959.

10. Yadav RN, Singh BK, Sharma SK, Sharma R, Soneja M, Sreenivas V, Myneedu VP, Hanif M, Kumar A, Sachdeva KS, Paramasivan CN, Vollepore B, Thakur R, Raizada N, Arora SK, Sinha S. 2013. Comparative evaluation of GenoType MTBDRplus line probe assay with solid culture method in early diagnosis of multidrug resistant tuberculosis (MDR-TB) at a tertiary carecentre in India. PLoS One 8:e72036.

11. Maningi NE, Malinga LA, Antiabong JF, Lekalakala RM, Mbelle NM. 2017. Comparison of line probe assay to BACTEC MGIT 960 system for susceptibility testing of first and second-line anti-tuberculosis drugs in a referral laboratory in South Africa. BMC InfectDis 17:795.

12. Bang D, Andersen SR, Vasiliauskienė E, Rasmussen EM. 2016. Performance of the GenoType MTBDRplus assay (v2.0) and a new extended GenoType MTBDRsl assay (v2.0) for the molecular detection of multi- and extensively drug-resistant Mycobacterium tuberculosis on isolates primarily from Lithuania. Diagn Microbiol Infect Dis 86: 377–381.

13. Heep M, Brandstätter B, Rieger U, Lehn N, Richter E, Rüsch-Gerdes S, Niemann S. 2001. Frequency of *rpoB*mutations inside and outside the cluster I region in rifampin-resistant clinical *Mycobacterium tuberculosis*isolates. J Clin Microbiol 39: 107–110.

14. Hillemann D, Weizenegger M, Kubica T, Richter E, Niemann S. 2005. Use of the genotype MTBDR assay for rapid detection of rifampin and isoniazid resistance in *Mycobacterium tuberculosis* complex isolates. J Clin Microbiol 43: 3699–3703.

15. Hillemann D, Rusch-Gerdes S, Richter E. 2007. Evaluation of the GenoType MTBDRplus assay for rifampin and isoniazid susceptibility testing of *Mycobacterium tuberculosis* strains and clinical specimens. J Clin Microbiol 45:2635–2640.

16. Felkel M, Exner R, Schleucher R, Lay H, Autenrieth IB, Kempf VA, Frick JS. 2013. Evaluation of Mycobacterium tuberculosis drug susceptibility in clinical specimens from Nigeria using genotype MTBDRplus and MTBDRsl assays. Eur J Microbiol Immunol (Bp) 3: 252–257.

17. Liu Q, Li GL, Chen C, Wang JM, Martinez L, Lu W, Zhu LM. 2017. Diagnostic Performance of the GenoType MTBDRplus and MTBDRsl Assays to Identify Tuberculosis Drug Resistance in Eastern China. Chin Med J (Engl) 130:1521–1528.

18. Ramaswamy SV, Reich R, Dou SJ, Jasperse L, Pan X, Wanger A, Quitugua T, Graviss EA. 2003. Single nucleotide polymorphisms in genes associated with isoniazid resistance in *Mycobacterium tuberculosis*. Antimicrob Agents Chemother 47:1241–1250.

19. van Soolingen D, de Haas PE, van Doorn HR, Kuijper E, Rinder H, Borgdorff MW. 2000. Mutations at amino acid position 315 of the *katG*gene are associated with high-level resistance to isoniazid, other drug resistance, and successful transmission of *Mycobacterium tuberculosis*in The Netherlands. J Infect Dis 182: 1788–1790.

20. Zhang Z, Lu J, Liu M, Wang Y, Qu G, Li H, Wang J, Pang Y, Liu C, Zhao Y. 2015. Genotyping and molecular characteristics of multidrug-resistant Mycobacterium tuberculosis isolates from China. J Infect 70:335–345.

21. Tolani MP, D’Souza DT, Mistry NF. 2012. Drug resistance mutations and heteroresistance detected using the GenoType MTBDRplus assay and their implication for treatment outcomes in patients from Mumbai, India. BMC Infect Dis 12:9.

22. Ramaswamy S, Musser JM. 1998. Molecular genetic basis of antimicrobial agent resistance in *Mycobacterium tuberculosis*: 1998 update. Tuber Lung Dis 79:3–29.

23. Somoskovi A, Parsons LM, Salfinger M. 2001. The molecular basis of resistance to isoniazid, rifampin, and pyrazinamide in *Mycobacterium tuberculosis*. Respir Res 2:164–168.

24. Brossier F, Veziris N, Aubry A, Jarlier V, Sougakoff W. 2010. Detection by GenoType MTBDRsl test of complex mechanisms of resistance to second-line drugs and ethambutol in multidrug-resistant *Mycobacterium tuberculosis* complex isolates. J Clin Microbiol 48: 1683–1689.

25. Hillemann D, Rüsch-Gerdes S, Richter E. 2009. Feasibility of the genotype MTBDRsl assay for fluoroquinolone, amikacin/capreomycin, and ethambutol resistance testing of mycobacterium tuberculosis strains and clinical specimens. J Clin Microbiol 47:1767–1772.

26. Kiet VS, Lan NT, An DD, Dung NH, Hoa DV, van Vinh Chau N, Chinh NT, Farrar J, Caws M. 2010. Evaluation of the MTBDRsl test for detection of second-line-drugresistance in *Mycobacterium tuberculosis*. J Clin Microbiol 48:2934–9.

27. Li J, Gao X, Luo T, Wu J, Sun G, Liu Q, Jiang Y, Zhang Y, Mei J, Gao Q. 2014. Association of *gyrA*/*B* mutations and resistance levels to fluoroquinolonesin clinical isolates of Mycobacterium tuberculosis. Emerg Microbes Infect 3:e19.

28. Zeng X, Jing W, Zhang Y, Duan H, Huang H, Chu N. 2017. Performance of the MTBDRsl line probe assay for rapid detection of resistance to second-line anti-tuberculosis drugs and ethambutol in China. Diagn Microbiol Infect Dis 89:112–117.

29. Lee YS, Lee BY, Jo KW, Shim TS. 2017. Performance of the GenoType MTBDRsl assay for the detection second-line anti-tuberculosis drug resistance. J Infect Chemother 23:820–825.

30. Ajbani K, Nikam C, Kazi M, Gray C, Boehme C, Balan K, Shetty A, Rodrigues C. 2012. Evaluation of Genotype MTBDRsl Assay to Detect Drug Resistance Associated with Fluoroquinolones, Aminoglycosides and Ethambutol on Clinical Sediments. PLoS One 7:e49433.

31. Maus CE1, Plikaytis BB, Shinnick TM. 2005. Molecular analysis of cross-resistanceto capreomycin, kanamycin, amikacin, and viomycin in *Mycobacterium tuberculosis*. Antimicrob Agents Chemother 49:3192–3197.

32. Via LE, Cho SN, Hwang S, Bang H, Park SK, Kang HS, Jeon D, Min SY, Oh T, Kim Y, Kim YM, Rajan V, Wong SY, Shamputa IC, Carroll M, Goldfeder L, Lee SA, Holland SM, Eum S, Lee H, Barry CE 3rd. 2010. Polymorphisms associated with resistance and cross-resistance to aminoglycosides and capreomycin in Mycobacterium tuberculosis isolates from South Korean patients with drug-resistant tuberculosis. J ClinMicrobiol 48:402–411.

33. Georghiou SB, Magana M, Garfein RS, Catanzaro DG, Catanzaro A, Rodwell TC. 2012. Evaluation of genetic mutations associated with Myeobacterium tuberculosis resistance toamikacin, kanamycin and capreomycin: a systematic review. PLoS One 7:e33275.

34. Cheng S, Cui Z, Li Y, Hu Z. 2014. Diagnostic accuracy of a molecular drug susceptibility testing method for the antituberculosis drug ethambutol: a systematic review and meta-analysis. J Clin Microbiol 52: 2913–2924.

35. Huang WL, Chi TL, Wu MH, Jou R. 2011. Performance assessment of the GenoType MTBDR*sl* test and DNA sequencing for detection of second-line and ethambutol drug resistance among patients infected with multidrug-resistant *Mycobacterium tuberculosis*. J Clin Microbiol 49: 2502–2508.

36. Van Deun A, Wright A, Zignol M, Weyer K, Rieder HL. 2011. Drug susceptibility testing proficiency in the network of supranational reference laboratories. Int J Tuberc Lung Dis 15: 116–124.

37. Crudu V, Stratan E, Romancenco E, Allerheiligen V, Hillemann A, Moraru N. 2012. First evaluation of an improved assay for molecular genetic detection of tuberculosis as well as rifampin and isoniazid resistances. J ClinMicrobiol 50: 1264–1269.

38. Tagliani E, Cabibbe AM, Miotto P, Borroni E, Toro JC, Mansjö M, Hoffner S, Hillemann D, Zalutskaya A, Skrahina A, Cirillo DM. 2015. Diagnostic Performance of the new version of GenoType MTBDR*sl* (V2.0) Assay for detection of resistance to Fluoroquinolones and Second-Line Injectable Drugs: a Multicenter study.J Clin Microbiol 53: 2961–2969.

39. HAIN LifeScience. 2015. GenoType MTBDRsl VER 2.0 instructions for use. Document IFU-317A-01. HAIN LifeScience, Nehren, Germany.

